# CHEER: hierarCHical taxonomic classification for viral mEtagEnomic data via deep leaRning

**DOI:** 10.1101/2020.03.26.009001

**Authors:** Jiayu Shang, Yanni Sun

## Abstract

The fast accumulation of viral metagenomic data has contributed significantly to new RNA virus discovery. However, the short read size, complex composition, and large data size can all make taxonomic analysis difficult. In particular, commonly used alignment-based methods are not ideal choices for detecting new viral species. In this work, we present a novel hierarchical classification model named CHEER, which can conduct read-level taxonomic classification from order to genus for new species. By combining k-mer embedding-based encoding, hierarchically organized CNNs, and carefully trained rejection layer, CHEER is able to assign correct taxonomic labels for reads from new species. We tested CHEER on both simulated and real sequencing data. The results show that CHEER can achieve higher accuracy than popular alignment-based and alignment-free taxonomic assignment tools. The source code, scripts, and pre-trained parameters for CHEER are available via GitHub: https://github.com/KennthShang/CHEER.

## 1 Introduction

Metagenomic sequencing, which allows us to directly obtain total genomic DNAs from host-associated and environmental samples, has led to important findings in many areas, such as digestive health [1]. While bacteria are the main focus of most metagenomic sequencing projects, there are fast accumulation of viral metagenomic data with sequencing viruses as the main purpose [2]. There are different types of viruses. We are mainly concerned with RNA viruses because many RNA viruses are notorious human pathogens, such as Influenza A, Human immunodeficiency virus (HIV), Ebola, SARS-CoV, and recently identified 2019-nCoV causing Wuhan pneumonia. Unlike DNA viruses, RNA viruses, which contain RNA genomes, lack faithful proofreading mechanisms during replication and thus can produce a group of related but different viral strains infecting the same host. This high genetic diversity within and across different hosts poses a great challenge for designing long-term protection strategies against these infectious diseases. For example, as the circulating strains can change every year, flu vaccine has to be administered every year.

Advances in viral metagenomics have contributed significantly to new RNA virus discovery. According to a survey by Woolhouse et. al, the number of newly identified RNA viruses is changing from 1,899 in 2005 to 5,561 in 2018, which is 3 times of increase [3]. Large-scale RNA virus sequencing projects using next-generation sequencing technologies have been conducted for different species. For example, Shi et al. have discovered a large number of new RNA viruses by sequencing samples from invertebrate and vertebrate animals [4, 5]. Claire et al. have discovered 25 new RNA viruses by sequencing samples associated with different drosophilid [6]. Bolduc et al. sequenced RNA viruses from Archaeal and bacterial samples [7]. Given fast accumulation of viral metagenomic data and expected discovery of new viruses, a key step is to conduct composition analysis for these data and assign taxonomic groups for possibly new species.

Composition analysis can be conducted at the read level or contig level. There is no doubt that contigs can provide more information than short reads in taxonomic classification. However, metagenomic assembly is still one of the most challenging computational problems in bioinformatics. Unlike single genome assembly, metagenomic assembly is more likely to produce chimeric contigs that can contain regions from different species. Thus, metagenomic assembly is often preceded by read binning that groups reads with the same or similar taxonomic labels [8] in order to achieve better performance.

In this work, we design a read level taxonomic classification tool (named as “CHEER”) for labeling new species in viral metagenomic data. To detect known species with available reference genomes, alignment-based methods can provide sufficient information for both classification and abundance estimation. This function has been incorporated in existing pipelines [9,10,11] and is thus not the focus of our tool. Instead, our tool is designed to handle the challenging cases of assigning taxonomic labels for reads of new species, which have not been observed before. This problem was clearly formulated in metagenomic phylogenetic classification tools such as Phymm and PhymmBL [8]. We first define the classification problem following the formulation in Phymm: the classification problem is the assignment of a phylogenetic group label to each read in input data sets. As we are only interested in the hard case of classifying reads of new species, which have not been sequenced and thus no label exists for these species, the expected labels for these reads are the higher-rank taxonomic groups such as genus, family, or order. All our data and experiments are designed so that the test species are masked in the reference database and the trained models.

### 1.1 Related Work

Previous experiments have shown that nucleotide-level homology search is not sensitive enough for assigning higher rank taxonomic groups to reads from new species [9, 10]. Protein-based homology search is thus preferred for phylogenetic classification [11, 12]. In particular, a recent tool, VirusSeeker [13] utilized BLASTx to classify reads into bacteria, phages, and other viruses. It is not clear whether the protein homology search is able to generate accurate classification at lower ranks. We will evaluate this in our experiments.

There are also alignment-free methods for phylogenetic classification. Many of those tools implemented different machine learning models and were mainly designed for bacteria. For example, RDP [14] adopted a Naïve Bayes based classifier that can automatically learn features from the reads and assigned taxonomic labels to sequences from bacterial and archaeal 16S genes and fungal 28S genes. The NBC classifier [15, 16] encoded metagenomic reads using k-mers and implemented a bag-of-word model to achieve the read-level taxonomic classification. Phymm [8] utilized interpolated Markov models (IMMs) for metagenomic read classification. These learning-based models enable users to label reads originating from new species.

The previous works have shown that sequence composition is still an important and informative feature for phylogenetic classification. Recent applications of deep learning models have achieved better performance than conventional machine learning methods in different sequence classification problems [17]. In particular, the convolution filters in Convolutional Neural Network models (CNN) can represent degenerated sequence motifs and thus help CNN models to learn abstract composition-based features. For example, frequently activated convolution filters learned by DeepFam often represented the conserved motifs in different protein domain families [18].

The successful applications of deep learning in sequence classification motivated us to design a novel deep learning-based classification model for assigning taxonomic groups for new species in viral metagenomic data.

Our main contributions are summarized below. First, our deep learning model can assign higher rank phylogenetic group labels (such as genus) for reads from new species. To achieve this goal, our model organizes multiple CNN-based classifiers in a tree for hierarchical taxonomic group assignment. Second, to enable our model to capture as much information as possible from short reads, we implemented and compared two encoding methods: one hot vs. embedding. Third, as viral metagenomic data usually contains contaminations from either the host genomes or other microbes, we formulated the pre-processing step as an open set problem in targeted image classification and rejected non-viral reads by choosing appropriate negative training sets. We tested our model on both simulated metagenomic data and also real sequencing data. The results show that our model competes favorably with other popular methods.

## 2 Method

In this section, we will introduce our method for viral metagenomic taxonomic classification. First, we will show the architecture of CHEER, which is a hierarchical classification model from order to genus. Second, classifiers at each level will be described. Third, we will introduce DNA sequence encoding using skip-gram based word embedding and compare it with one-hot encoding. Forth, we will describe the rejection layer, following the idea of the open set problem, which is adopted to filter reads not belonging to RNA viruses. Finally, we will detail our dataset used in training and validation.

### 2.1 Hierarchical classification model

The architecture of our model is sketched in Fig. 1. The key component is a tree model that consists of multiple classifiers from order to genus. In order to conduct phylogenetic classification for reads from new species, or even new genus, our classification is conducted using a top-down approach from the root to the leaf node. The top layer is a trained CNN that can reject reads not belonging to RNA viruses, which is an important preprocessing step for viral metagenomic data because of the contamination from the host genome or other microbes. Then, the filtered reads (i.e. mostly RNA viral reads) will be fed into the hierarchical classification model, which is referred to as the tree model hereafter. Specifically, the order-level classifier is responsible for classifying the reads into their originating orders. Then each order has a separately trained classifier for assigning input reads to families within that order (O1 to O3 in Fig. 1). For all the reads assigned into one family, the family classifier (F1-F7) will assign the reads to different genus within that family. As an example, a path for classifying a read from a species in genus 7 is highlighted in Fig. 1. CHEER also implemented early stop functions at each level for stopping the classification path at higher ranks. This function can accommodate taxonomic classification for species from new genus or even higher ranks, which is possible for RNA viruses. It is convenient to add fine-grained classifiers within each genus so that we can assign species-level labels. However, as our focus is to conduct phylogenetic classification for reads from new species that do not have a species labels in the training data, we did not include the species-level assignment in this work.

**Fig. 1.**
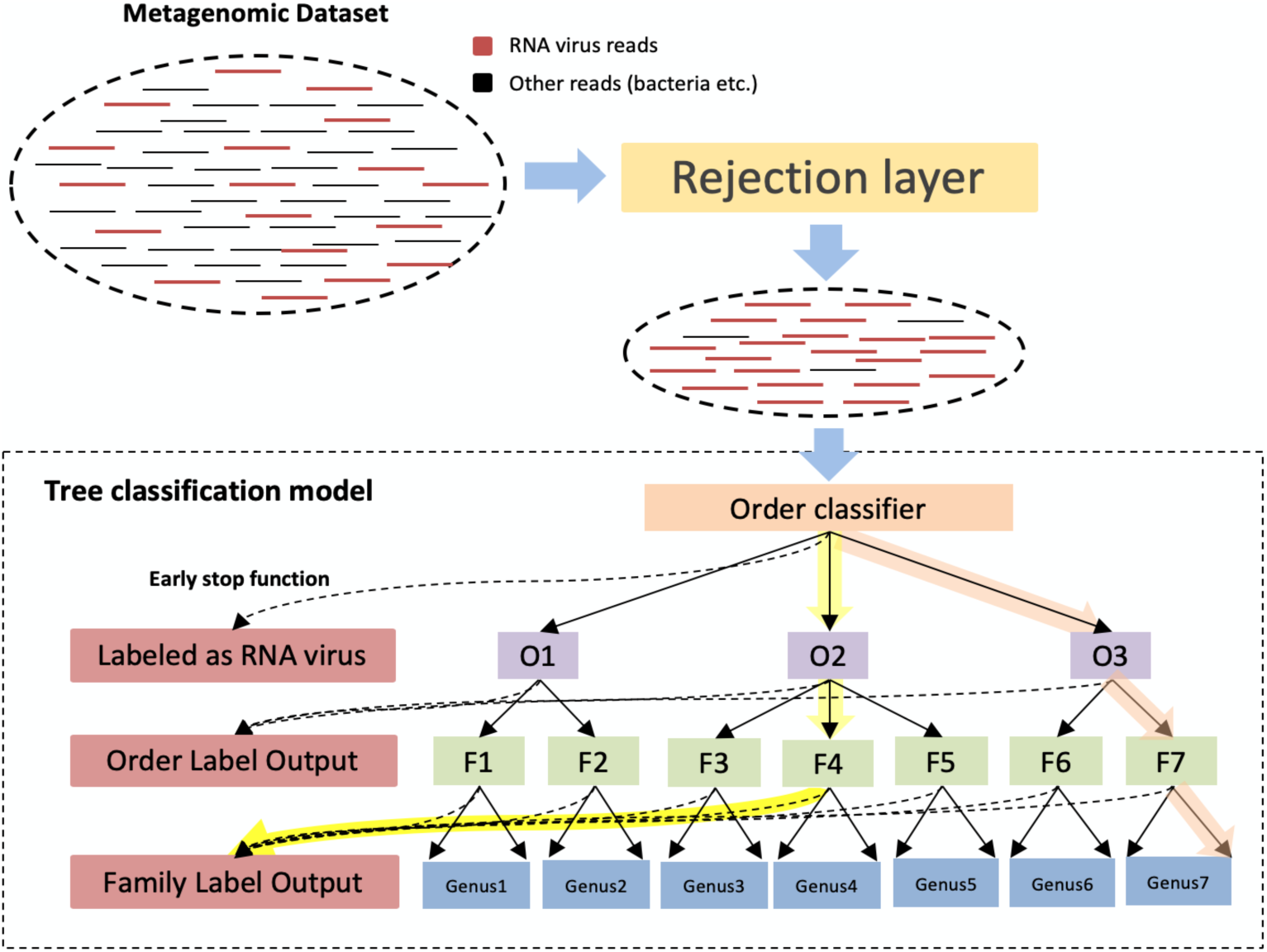
The flow chart and a simplified tree classification structure of CHEER. Order classifier: classify reads into orders O1 to O3. O1-O3: classifiers within O1, O2, and O3, which assign family-level labels within each order. F1-F7: classifiers within each family from F1 to F7, which assign genus-level labels within each family. The dash line under each classifier shows the early stop function. The highlighted path with orange color shows the top-down classification path for a read from genus 7. The highlighted path with yellow shows the top-down classification path that stops at the family level determined by the early stop function.

Alternative designs of hierarchical classification work, like [19, 20], either build a classifier for each taxonomic rank or build a binary classifier for each class. We implemented a structure that built just one classifier for each taxonomic rank and compared different structures’ performance. According to the result shown in [Supplementary file 1], the structure shown in Fig.1 is better than one classifier per rank. Although one classifier per rank requires fewer classifiers, it needs to train many more parameters because the number of labels for each classifier increases by times. As a result, we observe overfitting more frequently.

### 2.2 The structure of each classifier

Each classifier in the tree model is implemented using CNN, which has achieved superior performance in various sequence classification problems [18]. The convolution filters used in CNN resemble position-specific weight matrix of motifs in genomic sequences. Frequently activated convolution filters can represent well-conserved motifs among sequences in the same class. Thus, a large number of convolution filters will be used in our CNN in order to learn well-conserved sequence features in different classes. Deep CNNs in fields such as computer vision often have multiple convolution layers in order to obtain more abstract features from training images. Here, we use a wide convolution design rather than a deep one, following the idea in [18]. Fig. 2 shows the classifier with the default hyper-parameters. First, reads will be encoded in matrices as input to the convolution layer using two methods. We will describe and compare these two encoding methods later in the Section 2.3. Then, multiple convolution filters of different sizes are utilized to learn the conserved sequence features. According to our empirical study, we choose 3, 7, 11, 15 as the filter sizes, with 256 filters for each size respectively (see Fig. 2). Max pooling is then applied to each convolution filter’s output so that only the highest convolution value is kept. After the max-pooling layer, we have a 256-dimentional vector as the output of each convolution layer. We concatenate these four vectors into one 1024-dimentional vector and feed it to one fully connected layer with 512 hidden units. Dropout is applied to mitigate overfitting. The CrossEntropyLoss function supplied by Pytorch is adopted as an error estimation function. Due to the unbalanced size of each training class, we use the reciprocal of the proportion of each class as their error weight to help the model fairly learn from each class. And we utilize Adam optimizer with a 0.001 learning rate to update the parameters. The model is trained on HPCC with the 2080Ti GPU unit.

**Fig. 2.**
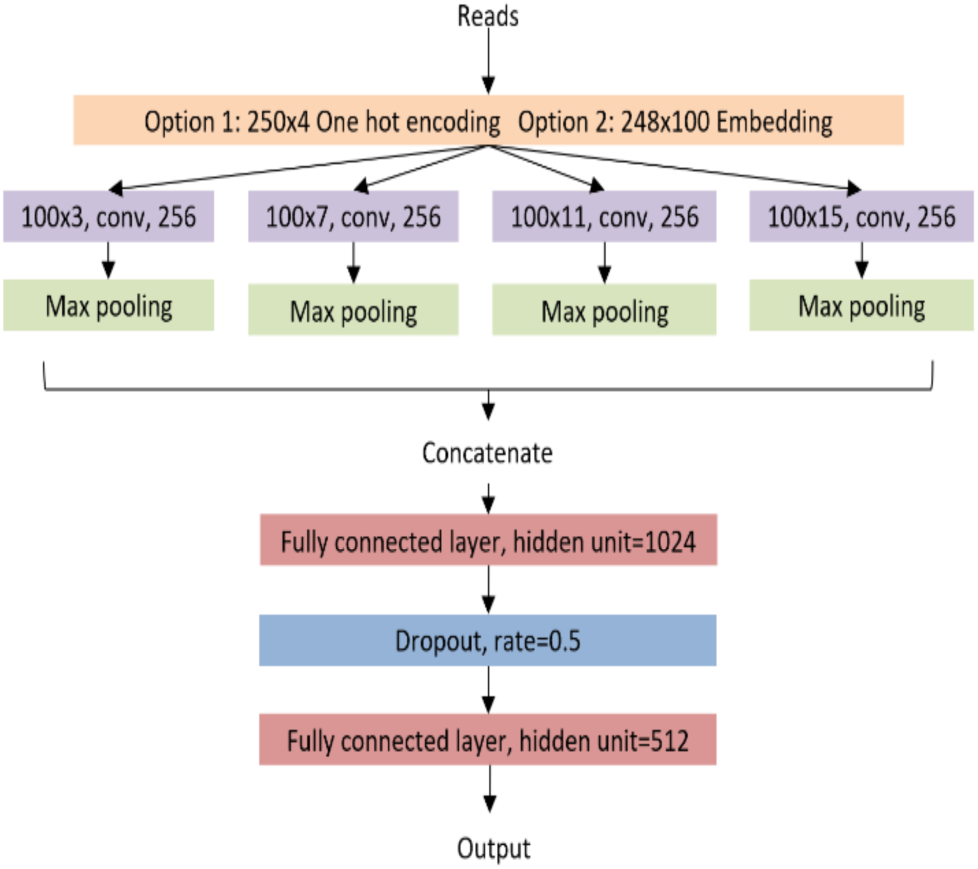
The CNN structure for each classifier in the tree model. As there are two options for read encoding, the figure only includes convolution filter size for k-mer embedding.

### 2.3 Read encoding

In previous deep learning-based sequence classification models [21, 22], one hot encoding was commonly adopted to convert a sequence into a matrix, which will then be used as input to convolution layers. There also exist some methods using k-mer composition [23] and frequency [15, 16] to encode the DNA sequence, similar to the bag-of-words model in the field of natural language processing. However, these k-mers representation cannot give complete sequential information about the original reads.

To incorporate both the k-mer composition and also their ordering information, we build a Skip-Gram [24] based embedding layer that can learn which k-mers tend to occur close to each other. Embedding is widely used in natural language processing to learn the semantic and syntactic relationships from sentences. A neural network with one hidden layer is trained to map a word to an n-dimensional vector so that the words that usually occur together will be closer in the n-dimension space. For our DNA read classification task, k-mers are the words and the embedding layer will map proximate k-mers into vectors of high similarity.

The main hyper-parameter when training the skip-gram model is the size of the training context *m*, which defines the maximum context location at which the furthest k-mer is taken for training. Specifically, for a k-mer at position *i* as the input to the skip-gram model, the output k-mers are its neighbors located at *i*+*jk*, where −m ≤ *j* ≤ m. Larger *m* results in more training samples with a cost of training time. In our model, the default value of *m* is 1. As shown in Fig. 3, we will sample 3 k-mers as our dataset at a time with the middle one as the input and the surrounding two as the output.

**Fig. 3.**
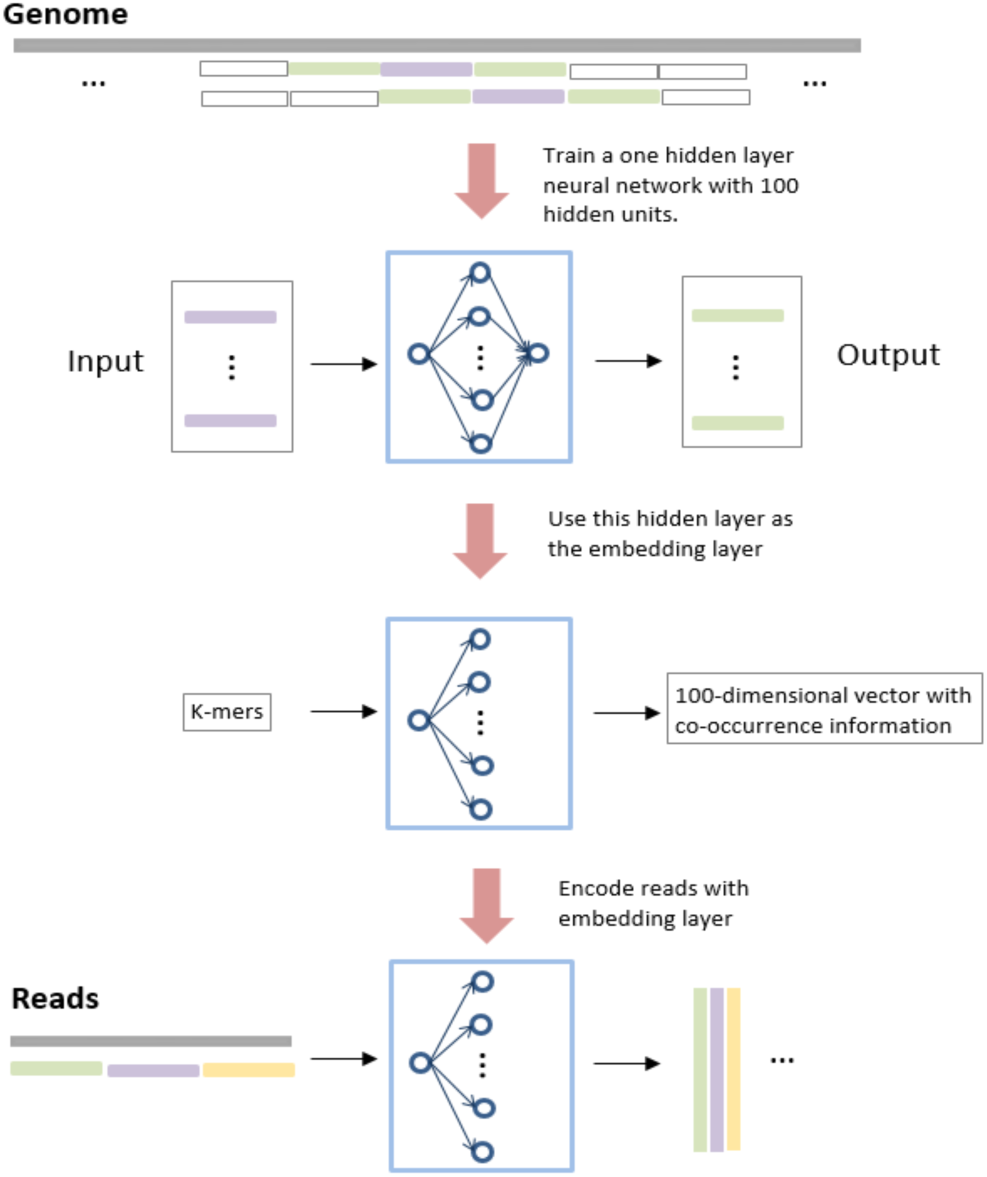
The training procedure of the embedding layer. There are 100 hidden units in the model.

As proven in [24], Skip-Gram is able to automatically learn the relationship between proximate words and thus is expected to help learn the order between k-mers in genomic data. We implemented a hidden layer with 100 hidden units shown in Fig. 3. After training the model, the co-occurrence features are embedded in the output of the hidden layer. Thus, a k-mer in a read will be represented as a 100-dimension vector, where each element in the vector is the output of the hidden unit in the hidden layer.

When we use this pre-trained embedding layer to encode the reads, we first split each read into sequential k-mers. Second, each k-mer is fed to the embedding layer and the output is a 100-dimension vector. Third, all these vectors are combined into a large matrix according to the original position of each k-mers. Finally, this matrix will be fed to the convolutional layer.

Based on our large-scale experiments, we found that classifiers are easy to overfit with increase of k. Thus, we choose 100-dimensional feature vectors with k being 3 as our default embedding layer parameter set based on our empirical analysis.

### 2.4 Viral read screening based on Open Set Problem

Due to the small genome size and limitations in library construction, it is not rare that many viral metagenomic datasets are still contaminated with the genomes of the host and other microbes. Viral read screening is thus a key pre-processing step before the downstream composition analysis. Specifically, our tree model should only be applied to viral reads. However, the close set nature of the classifier forces it to assign a known label to each input without being able to distinguish viral reads from others. In this work, we will reject non-viral reads by formulating it as an open set problem in targeted image classification [25].

In a majority of neural network models [26, 27], the logit of the last layer will be fed into the SoftMax function, which takes a vector of *n* real numbers as input and output a normalized probability vector for all labels/classes. However, for an irrelevant input from an unknown clade, all classes in the model tend to have low probabilities and thus applying a threshold on uncertainty can be used to reject unknown classes [25]. Along with this idea, serval approaches [28, 29] have been proposed to solve the open set problem.

In this work, we incorporate the pre-processing step in our hierarchical classification model by integrating a trained CNN and a SoftMax threshold. Contaminations such as bacteria will be rejected using empirically chosen SoftMax threshold τ. Our experiments show that SoftMax outputs a nearly uniform probability distribution of all labels if the input is not relevant. Thus, the SoftMax threshold τ is used to reject reads not belonging to either DNA or RNA viruses. However, using this thresholding method is not able to distinguish DNA viral reads from RNA viral reads due to their similar sequence composition. To accommodate this challenge, we designed a rejection classifier in the top layer using DNA viral reads as the negative class in the training (see Fig. 1).

### 2.5 The early stop function in the hierarchical classification

As the focus of our work is to assign labels for new species, the hierarchical classification ends at genus-level assignment, which is conducted by the bottom layer in Fig. 1. Given the fast accumulation of viral metagenomic data, there are cases where a new species does not belong to any existing genus and thus the label assignment should stop at the higher ranks. Thus, we allow our model to stop at a higher rank in the tree during the top-down classification. Towards this goal, we still apply SoftMax threshold in our early stop function, which stops reads from further classification by the child classifiers. This early stop function also serves as a quality control measure for preventing wrong classifications if the confidence is low. For example, when a read arrives, the model may be able to identify its order and family labels with high reliability but cannot decide which genus it belongs to. In this case, the model should only assign order and family label to this read rather than classifying it into a potentially wrong genus. To achieve this goal, SoftMax threshold is applied for every classifier in the tree. Only reads with a score higher than the threshold will be assigned to the child class.

An example of early stop is highlighted by the yellow path in Fig. 1. The model can classify the read into order 2 (O2) and then family 4 (F4) with high reliability but cannot decide which genus it belongs to.

### 2.6 Training and validation datasets

All the viruses used in the experiments were obtained from the RefSeq in the NCBI virus database. The viruses’ taxonomic classification was downloaded from Virus Taxonomy 2018b Release [30] published by the International Committee on Taxonomic of Virus (ICVT). Because training deep learning classifiers required enough samples, we removed the orders with only one family, families with only one genus, and genera with less than 3 species from our dataset. Finally, there are 6 orders, 23 families, and 55 genera remained. The RefSeq genomes are available at the following URL:https://www.ncbi.nlm.nih.gov/labs/virus/vssi/#/find-data/virus. The information of the taxa can be found in [Supplementary file 1].

As CHEER conducts read-level taxonomic classification, the training is also conducted using reads. In order to train our model on each complete virus genome, we generated simulated reads by uniformly extracting substrings of 250 bp with 200bp overlaps. All these reads formed the training set so that the model can learn the features from the whole genome. The validation set, however, is generated by read simulation tool WgSim [31] with specified error rate so that we can have a better estimation of the model’s performance on real sequencing data.

Some existing deep learning tools for taxonomic classification, like [21, 23], generate reads first, then randomly choose some of them as training set and others as testing set. This strategy cannot guarantee that the training and testing data sets have no overlaps from the same species. In order to test whether our model can learn the label of new species, we use test species masking in the training set. Specifically, we split our RNA virus database into two independent sets. For any test species, their reads are not used in the training data. Thus, the model will only use the knowledge of known species for the new species classification.

## 3 Results and Discussion

In this section, we will first give details of the hyperparameters of CHEER. Then we present the results of applying CHEER to both simulated and mock sequencing data. For the simulated data by WgSim, we evaluated how sequencing error rate affects the performance of CHEER and compared the taxonomic classification results with state-of-the-art. For the mock data, we tested CHEER’s performance in both taxonomic classification and its ability in distinguishing viral reads from others.

### 3.1 Training and Testing procedure

In our experiments, the minimum number of species within a genus is 3. We thus constructed three test sets, each containing reads from one species that is randomly chosen from each genus. The three test sets contain three different species, respectively. In total, 165 species from 55 genera are chosen to be “new” species for testing and the information of these species can be found in [Supplementary file 1].

We used WgSim to simulate reads from testing species. In order to evaluate the impact of sequencing error on CHEER, we simulated reads with an error rate of 0, 0.01, 0.02, and 0.05, respectively. For each classifier (CNN), we computed the classification accuracy, which is defined in equation (1). To ensure that the computation of the accuracy is not affected by the number of reads from different species, we constructed a balanced test read set by simulating the same number of reads for each test species (1,000 reads for each).

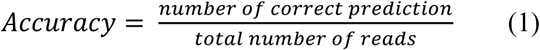

For each component classifier in the tree model, we record the average accuracy for multiple test species. For each taxa level, we calculate the average accuracy of all the classifiers in the same level.

### 3.2 Classification performance at each level

We present the results of CHEER at each level in this section. Because CHEER is designed for new species, we benchmark it against alignment-free models. Based on the results shown by Wood et al. [7], Naïve Bayes Classifier (NBC) outperformed other alignment-free models. Thus, we compared the performance of CHEER and NBC. NBC in [14] was designed for bacteria taxonomic classification and thus, we retrained NBC on our RNA virus dataset. We implemented the NBC model following [14] with the same parameters.

Fig. 4 shows the classification accuracy for genera. Since each family has a separately trained classifier, which conducts genus classification within that family, there are totally 14 classifiers at this level. Fig. 5 shows the accuracy for labeling families. There are totally 6 classifiers that assign reads to families within each of the 6 orders. Fig. 6 shows the mean accuracy at each rank. The family and genus level accuracy are the average accuracy of all classifiers in Fig. 4 and Fig. 5. There is only one classifier for order level classification.

**Fig. 4.**
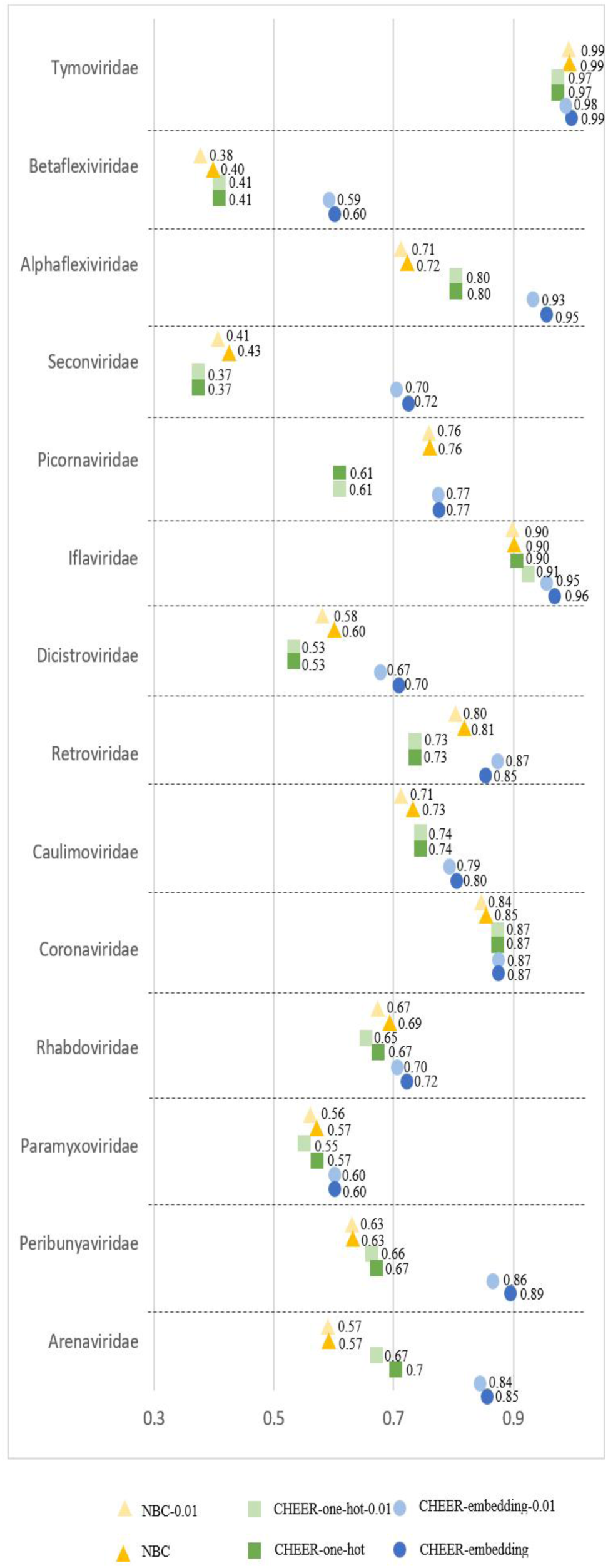
The classification accuracy at genus level. Y-axis: classifier within each family. X-axis: the accuracy. NBC: Naïve Bayes Classifier. CHEER-one-hot: one hot encoding option. CHEER: embedding layer option. Model name which suffix 0.01: testing reads with a sequencing error rate of 0.01.

**Fig. 5.**
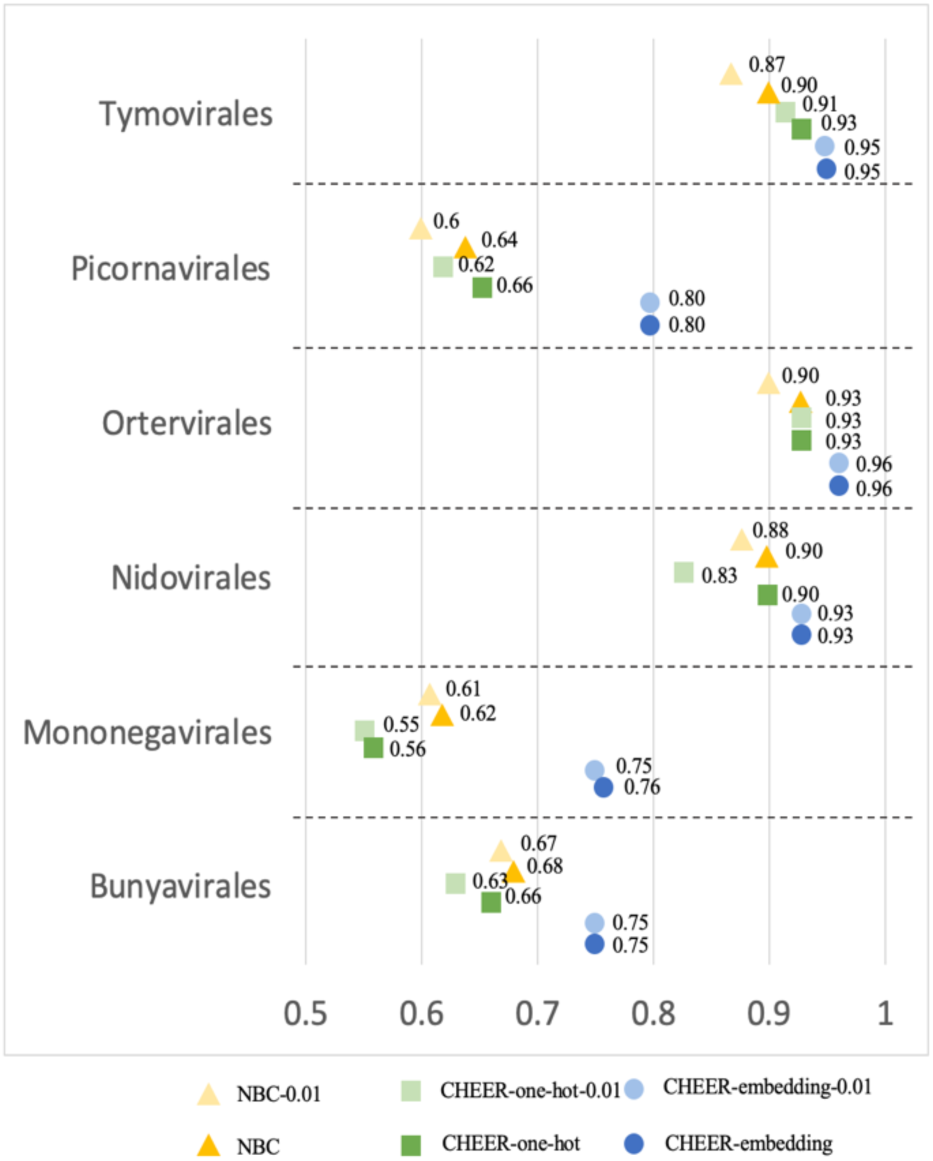
The classification accuracy at family level. Y-axis: the classifier within each order. X-axis: the accuracy. Refer to Fig. 4 for the meaning of the labels.

**Fig. 6.**
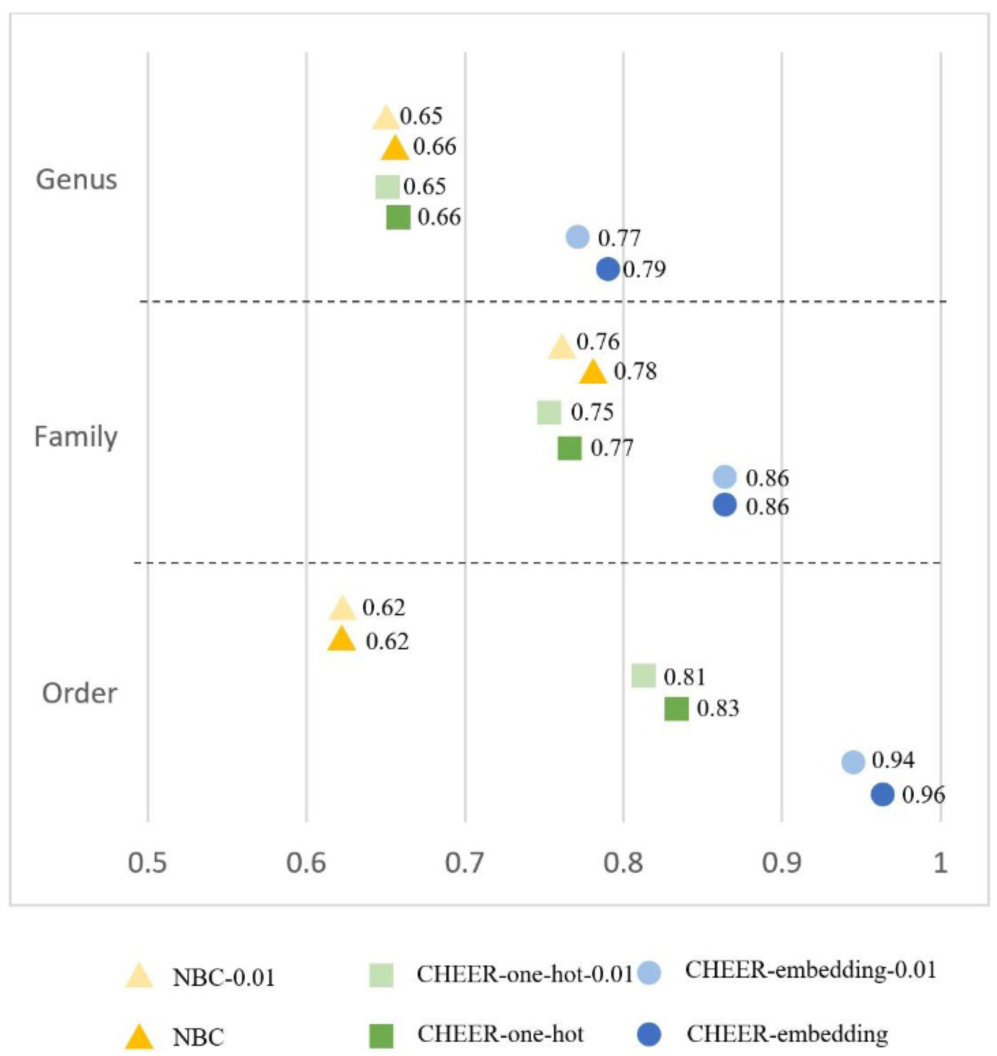
The mean accuracy at each level. Y-axis: the taxonomic rank. X-axis: the accuracy. Refer to Fig. 4 for the meaning of the labels.

As shown in Fig. 4, Fig. 5, and Fig. 6, CHEER has better performance than NBC across different ranks. We also compared the performance of using one-hot encoding and embedding layer in CHEER. The performance comparison at each rank shows that for most of the classifiers, the embedding layer improves the learning ability.

Fig. 4, Fig. 5, and Fig. 6 also revealed the robustness of CHEER. Although the classification accuracy decreases as the error rate increases, the decrease is smaller than 2%. The accuracy is almost identical for many classifiers. Thus, CHEER is not very sensitive to sequencing errors. We also conducted experiments for reads with higher error rates (0.02 and 0.05). The results can be found in [Supplementary file 1].

### 3.3 Real sequencing data

The next experiment is to evaluate our model on a real sequencing dataset and test CHEER’s performance in both taxonomic classification and non-viral read rejection.

We built a mock dataset by mixing Ebolavirus reads with gut bacterial reads from a gut amplicon sequencing dataset, which is chosen to ensure that we only use reads from bacteria. The taxonomic classification of Zaire Ebolavirus in the tree model is highlighted in Fig. 7. We chose one of the Ebola species named Zaire Ebolavirus as our testing species because it is the most dangerous of the five Ebolavirus species and was responsible for the 2014 West African Ebola outbreak. We downloaded the dataset from NCBI with accession number SRR1930021. These reads were sequenced by Illumina using whole-genome amplification strategy. Since Zaire Ebolavirus usually causes fatal hemorrhagic fever and abdominal pain, it is possible that this Ebola virus occurs with other gut bacteria. Thus, we created a mock viral metagenomic data by mixing these Ebola reads with a gut bacteria amplicon sequencing data in order to test whether CHEER can reject the bacterial reads at the top rejection layer. The gut bacteria dataset is downloaded from NCBI with accession number: SRR10714026. There are totally 2,737 Ebola virus reads and 51,280 gut bacterial reads in the mock data. After the rejection layer, the Ebola reads will be used to evaluate our tree model.

**Fig. 7.**
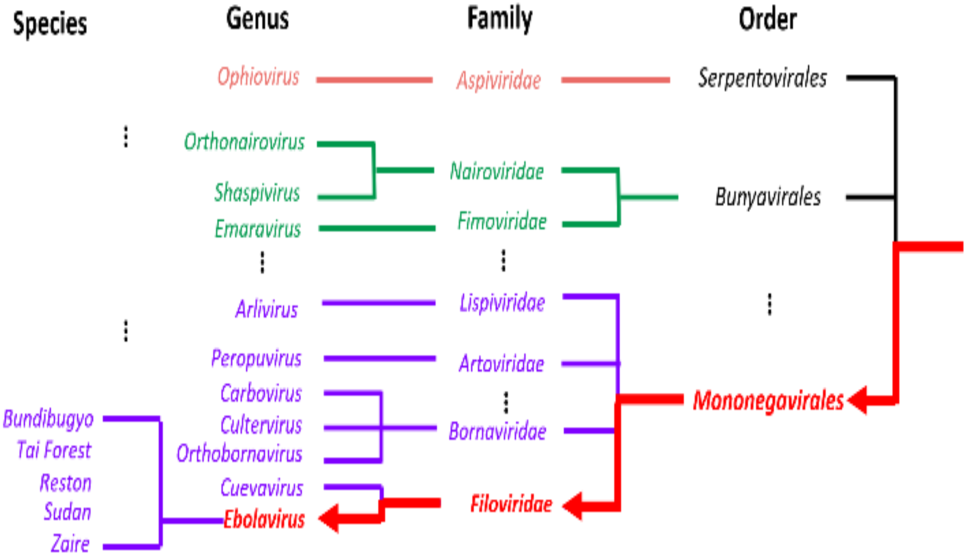
Taxonomy of Ebola. The red lines with arrows is the classification path for a new species in Ebolavirus in the tree model.

To treat Zaire Ebolavirus as a “new” species, we removed its genome from our training set and re-trained the whole pipeline. As shown in Fig. 7, after removing this species, we still have four Ebola species in our training set. However, our read mapping experiment using Bowtie2 [32] showed that none of the reads from Zaire Ebolavirus can be mapped to other Ebola species with the default parameters, indicating that using usual alignment-based methods will have difficulty to conduct genus-level assignment.

In our work, CHEER will first use rejection layer to remove the reads not belonging to the RNA virus. Second, the “Order classifier” will be applied to evaluate how many reads can be classified into Mononegavirales order. Third, “Mononegavirales classifier” takes all these Mononegavirales reads as input to predict how many reads will be assigned to the Filoviridae family. Finally, “Filoviridae classifier” is adopted to check how many reads belong to the Ebola genus.

#### 3.3.1 Evaluation of the rejection layer

Fig. 8 show the ROC curve of the rejection layer. True positive rate reveals how many Ebolavirus reads pass the rejection layer. The false positive rate reveals how many gut bacterial reads pass the rejection layer. We generated thresholds from 0 to 1 with step size 0.01 and used each threshold for the rejection layer. Then for each threshold, we recorded the true positive rate and false positive rate for plotting the ROC curve. As shown in Fig. 8, with the increase of the threshold, more bacterial reads are filtered by the rejection layer, at the expense of missing more real Ebola reads. The default threshold for the rejection layer is 0.6 in CHEER. User can adjust the threshold according to their needs.

**Fig. 8.**
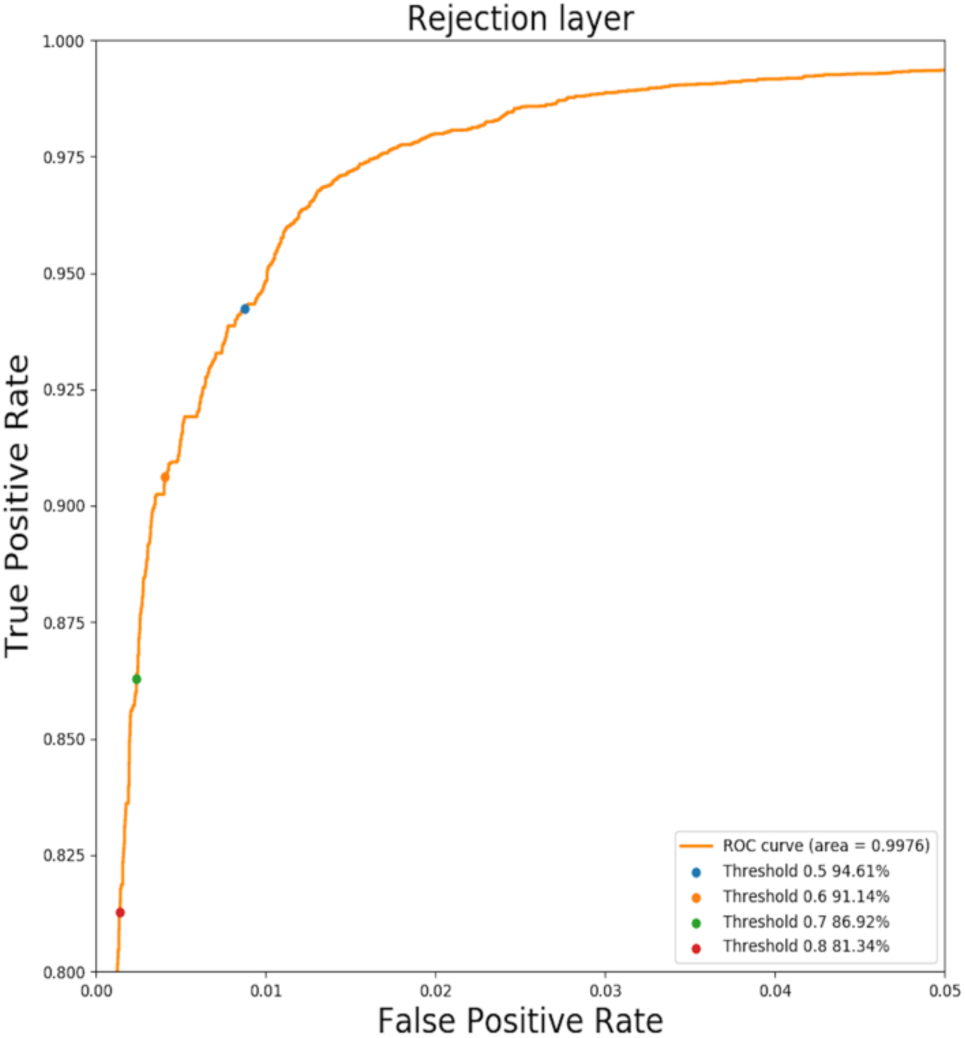
The ROC curve of the rejection layer for recognizing reads from Zaire Ebolavirus in the mock dataset. Y-axis: True positive rate of rejection layer. X-axis: False Positive rate of rejection layer. Four points corresponding to four thresholds are highlighted in the ROC curve. The percentage after each threshold means how many Ebolavirus reads pass the rejection layer.

#### 3.3.2 Comparison to the NBC model

After evaluating the rejection layer, we continued to test the hierarchical classification performance at each rank. We first compared our tree model with NBC. Then we also benchmarked CHEER with the method using protein-level alignment in Section 3.3.3.

The performance of the complete hierarchical classification pipeline of NBC and CHEER is shown in Fig. 9. From order to genus, each bar shows the accuracy of correctly classified reads by the parent classifier. For example, NBC correctly classified 1580 at the order level. Then, 1564 out of the 1580 reads are correctly labeled at the family level. While Fig. 4, 5, and 6 have shown the performance of CHEER and NBC at each taxonomic rank using the same set of reads as input for each level of classifiers, this experiment focused on comparing the hierarchical classification pipeline using only correctly predicted reads by the parent classifier. The results demonstrated that CHEER can classify Ebola reads with better performance. NBC can only classify 1,564 (57.1%) reads into Ebolavirus. CHEER, however, had totally 2407 (87.9%) reads classified correctly.

**Fig. 9.**
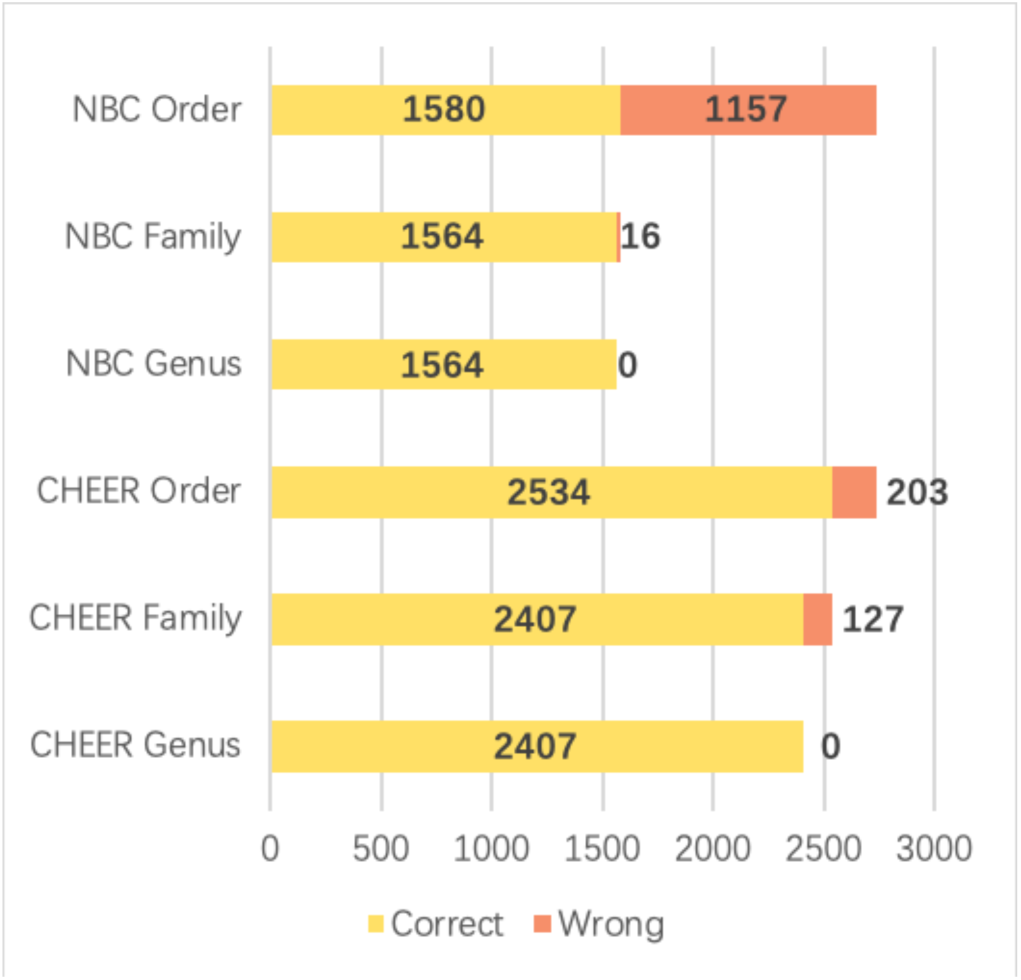
Zaire Ebolavirus read prediction at each rank using the top-down approach. Y-axis: different taxonomic rank with NBC and CHEER. X-axis: number of reads. Correct: reads with correct classification. Wrong: reads with wrong classification. Following the classification tree, only correct predictions (yellow bar) will be fed to the child classifier.

#### 3.3.3 Comparison with alignment-based methods

Although nucleotide-level alignment models are not ideal choices for predicting new species, as mentioned in [8], protein-level alignment such as BLASTx is still frequently used for homology search because of the conservation among homologous protein sequences. We followed VirusSeeker [10], which is a BLASTx based model, to align translated reads to virus-only protein database (VIRUSDB NR) provided by NCBI. Then, we kept alignments with E-value cutoff 0.001, which is used in VirusSeeker. The output of BLASTx shows that a read can often align to multiple proteins, making the phylogenetic classification ambiguous (see Fig. 11). Also, existing tools such as VirusSeeker often use the best alignment only, which does not always yield the highest accuracy. To optimize the usage of the alignment-based method, we implemented k-Nearest Neighbor (kNN) algorithm to record the best k alignments rather than only using the best one. The label of the input read is determined by the majority vote in the labels of the top k alignments.

**Fig. 10.**
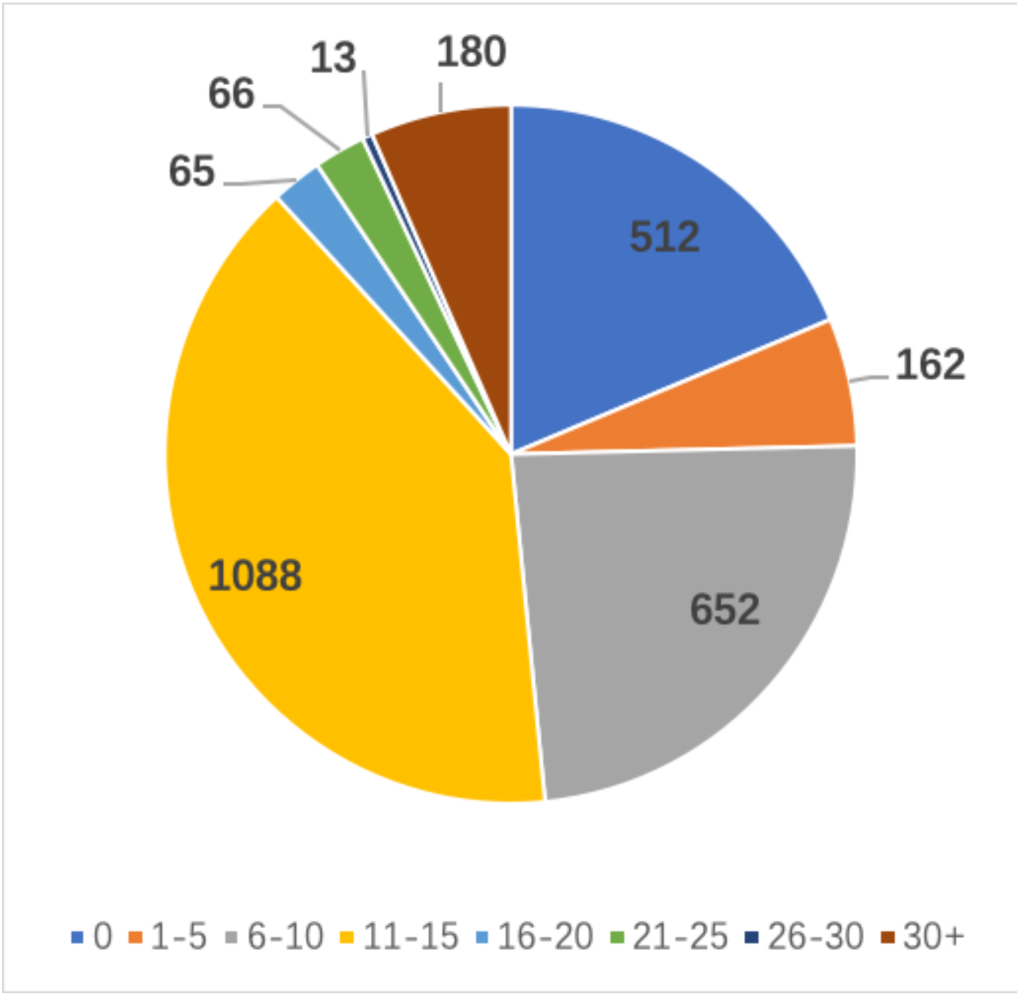
Distribution of BLASTx hit numbers with E-value threshold 0.001

**Fig. 11.**
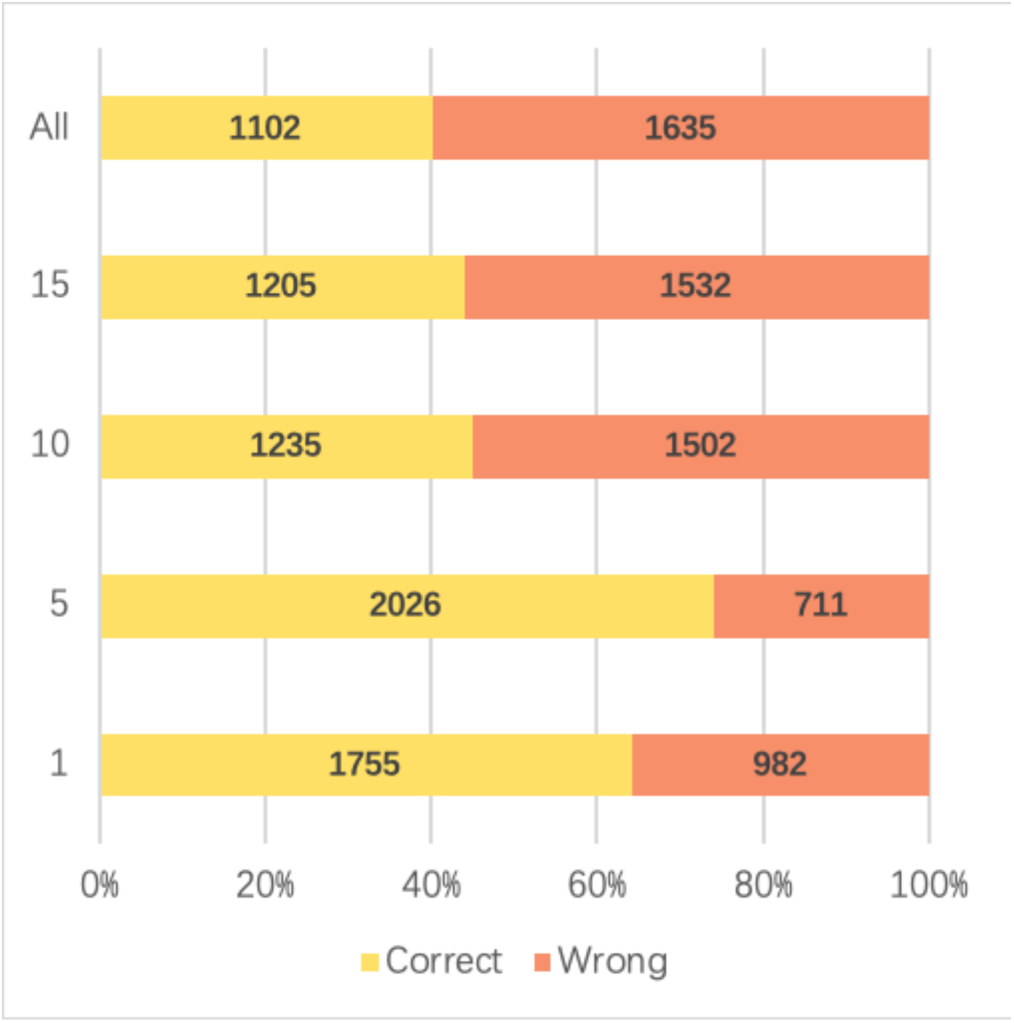
Ebola kNN BLASTx classification result with different k value. Y-axis: k value. X-axis: percentage of correct and wrong classification. “ALL”: all hits satisfying the E-value cutoff are used to run the majority voting program.

According to Fig. 10, most of the reads have hit numbers between 11 to 15 and 6 to10. Thus, we chose 1, 5, 10, 15 as our k when running kNN. There are also 512 reads that cannot be aligned with any VIRUSDB protein sequence. All these reads will be regarded as wrong prediction.

Fig. 11 shows the classification results of kNN. When we use the best hit as the prediction the accuracy is only 64%. The best performance of kNN will be 74% with 2,026 correct predictions when k is 5. But the accuracy sharply decreases if k is larger than 5.

Fig. 10 and Fig. 11 also revealed that it is hard for users to choose a proper k in real classification tasks. The most common value of the hit number is 11-15 shown in Fig. 11. The best kNN result, however, has a k equal to 5. Thus, comparing to this alignment-based method, CHEER, with 2,407 (87.9%) reads classified correctly, is more practical and reliable.

#### 3.3.4 Evaluation of the early stop function

The two main purposes of the early stop function are: 1) to handle the cases of a new species from an unseen genus or higher ranks; 2) to serve as a quality control measure for preventing wrong classifications in child classifiers. As Fig. 6 shows, the higher rank classification tends to be more accurate than lower ranks. Because the genus of Ebolavirus exists in our classification tree, the evaluation of the early stop function focuses on the second purpose.

Fig. 12 shows the percentage of how many reads are classified into Ebolavirus genus correctly for different classifiers. By allowing early stop, the number of wrong predictions decreased sharply. In short, although there are fewer reads being labelled at the genus level, the total number of reads with wrong classification is decreased. Thus, this is still a useful feature provided by CHEER.

**Fig. 12.**
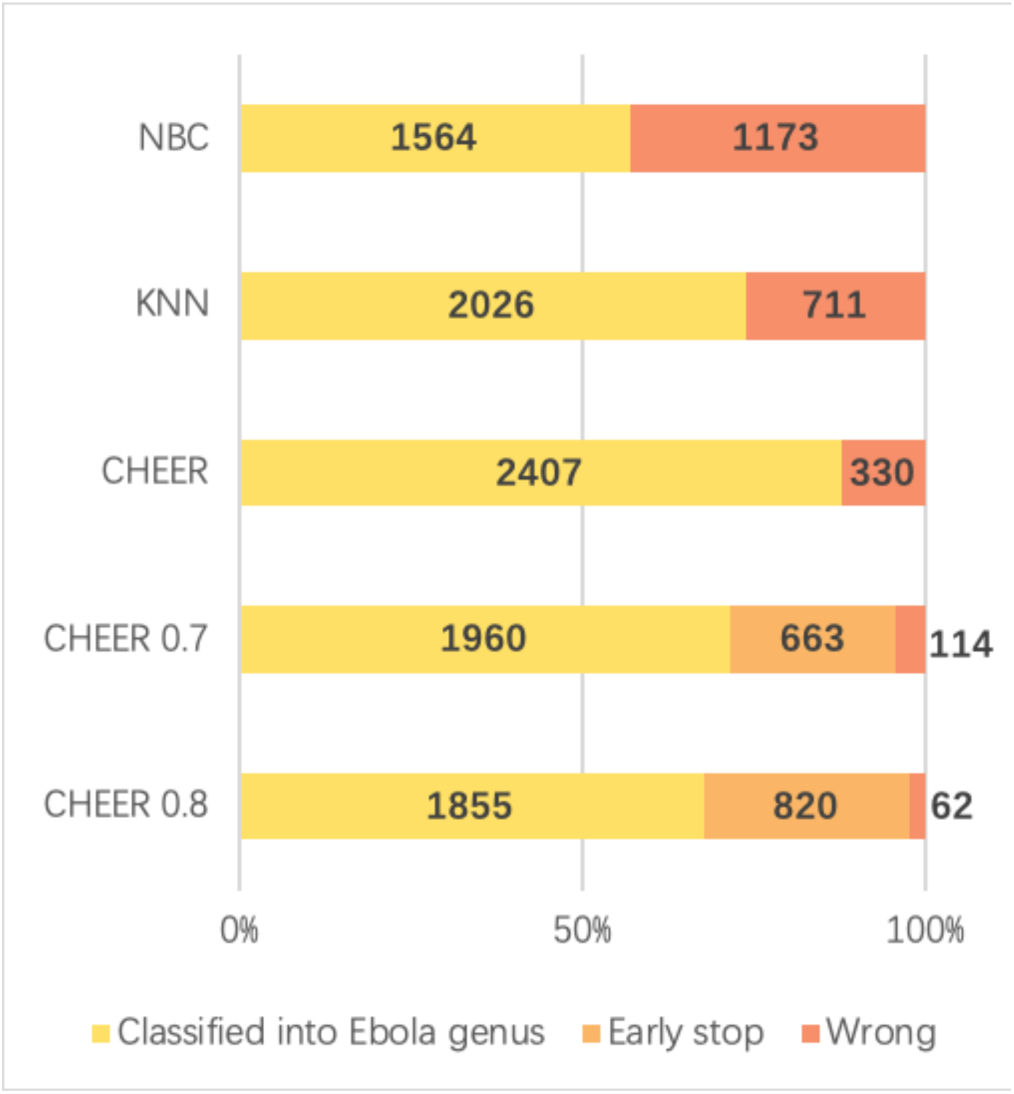
Comparison between different models. Y-axis: name of different classifier. NBC: Naïve Bayes classifier. kNN: k-Nearest Neighbor model. CHEER: without early stop threshold. CHEER 0.7: early stop threshold 0.7. CHEER 0.8: early stop threshold 0.8. X-axis: percentage and number of correct, wrong, and early stopped classifications. “Classified into Ebola genus (yellow)”: correctly classified reads. “Early stop (light orange)”: reads that are stopped at higher ranks. “Wrong (dark orange)”: misclassified reads.

#### 3.3.5 Comparison of execution speed

Another significant advantage of CHEER is the high classification speed. Although training the model uses heavy computing resources, the inference procedure of this algorithm is fast. And with the help of GPU acceleration, the training time also decreases a lot.

In the experiments shown in Section 3.4, it takes around 46 minutes to run BLASTx against the virus protein database. Our method, however, only takes 7 second for the preprocessing step to convert the raw sequence into a vector, and then takes less than 35 second to make the prediction for all 2,737 reads.

## 4 Conclusion

In this work, all training data are RNA virus genomes downloaded from the NCBI RefSeq database. The training reads are extracted uniformly from the genome to make sure that classifiers can learn the features from the whole genome. To evaluate the model on new species detection, all species in the test set are excluded from the training data. Error rates are also added to evaluate the robustness of the model. The results reveal that the model is tolerant to the sequencing error. In addition, a SoftMax threshold rejection layer is applied to filter reads from non-RNA viruses. The SoftMax threshold in each classifier will also help early stop classification with low confidence and thus decreases the wrong predication rate. We show a case study on real-world data to identify how this threshold will influence the rejection classifier and early stop function. The results show that CHEER competes favorably with state-of-the-art tools.

## Acknowledgement

The authors would like to acknowledge the computational support from the high-performance computing system at Electrical Engineering Department of City University of Hong Kong.

## Funding

this work was supported by the City University of Hong Kong Project 7200620 and Hong Kong RGC GRF grant 9042828.

